## Supplementary figures and images for "Non-allometric expansion and enhanced compartmentalization of Purkinje cell dendrites in the human cerebellum"

### Supplemental Figure 1

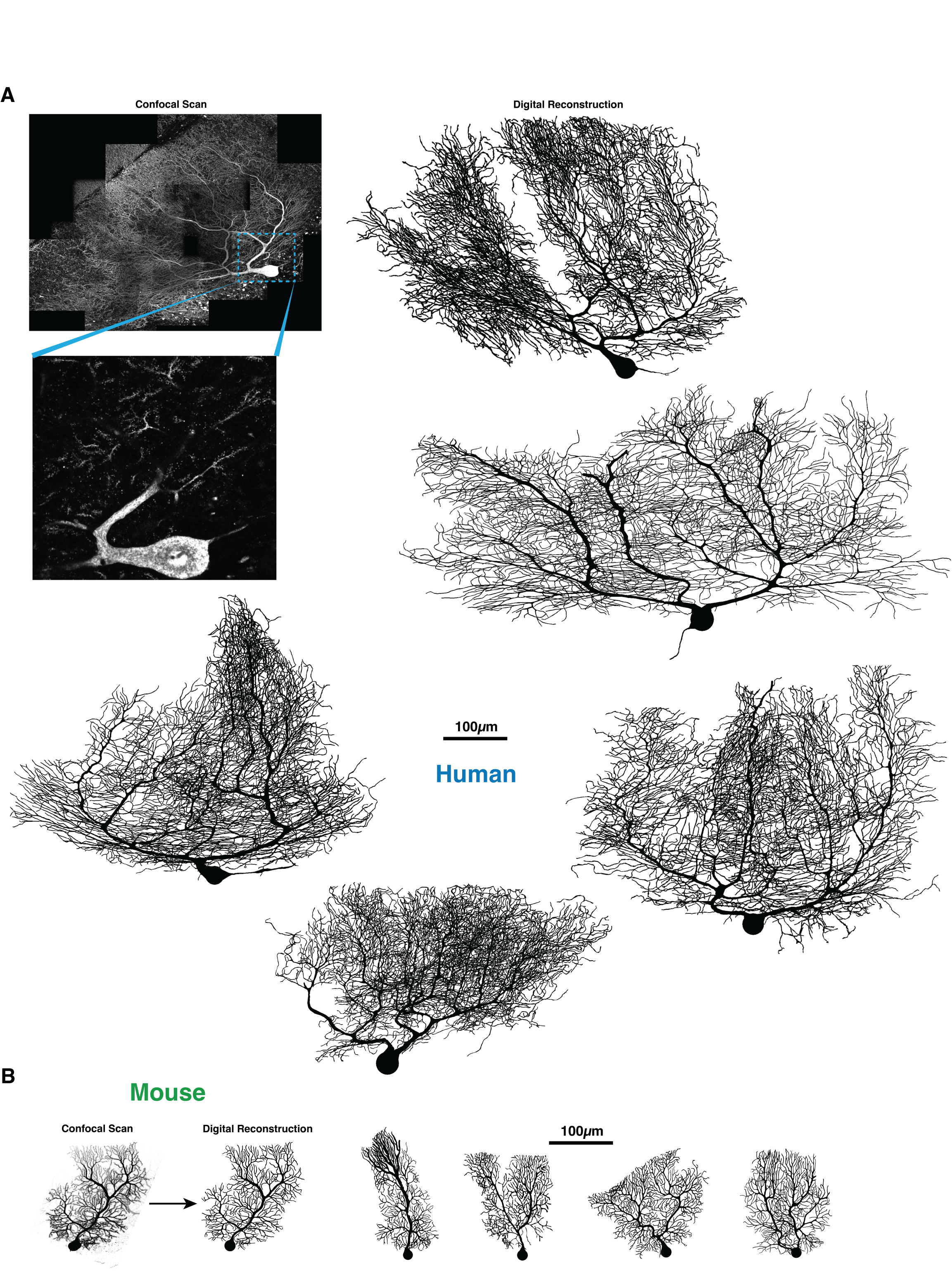

### Supplemental Figure 2

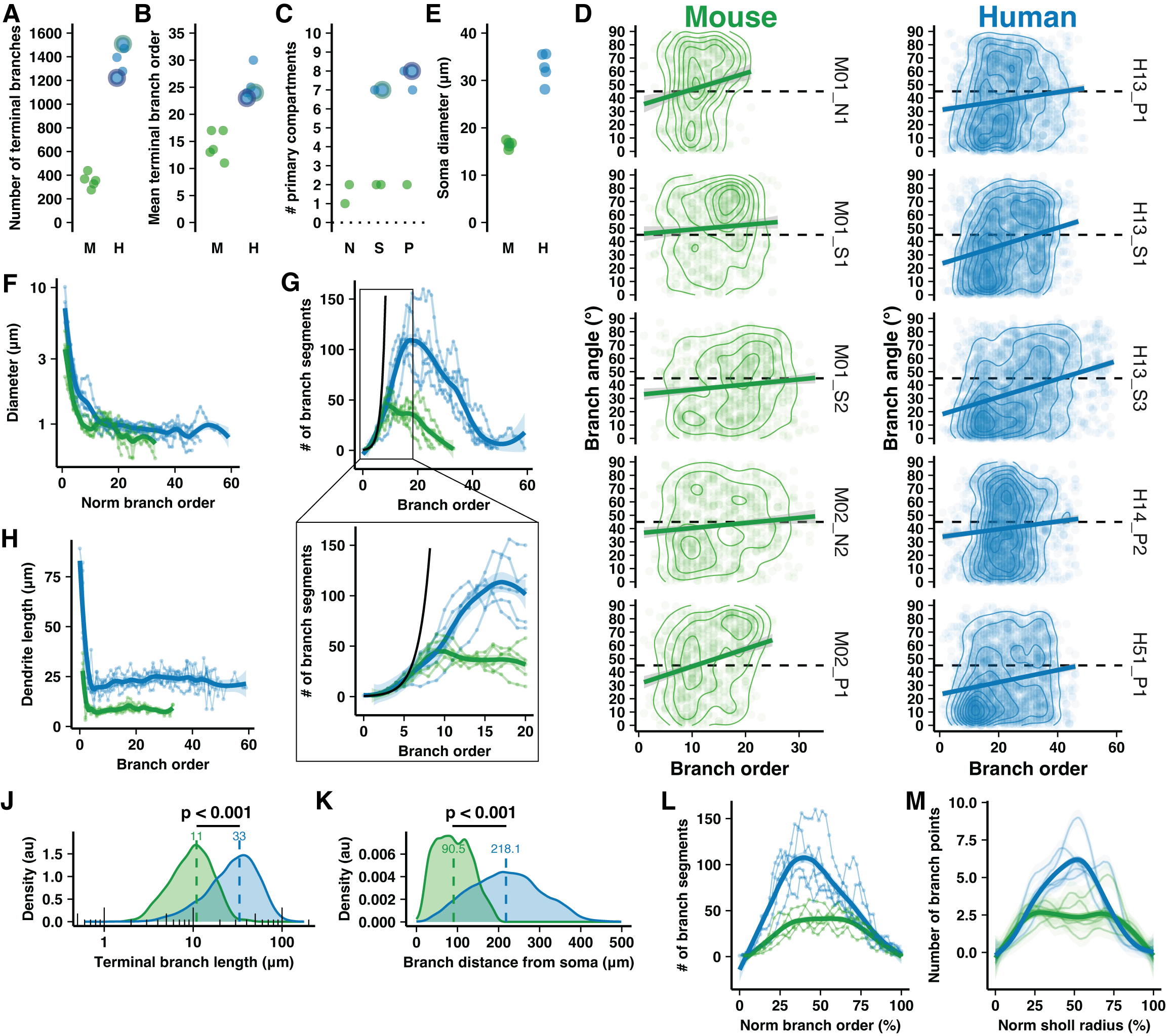

### Supplemental Figure 3

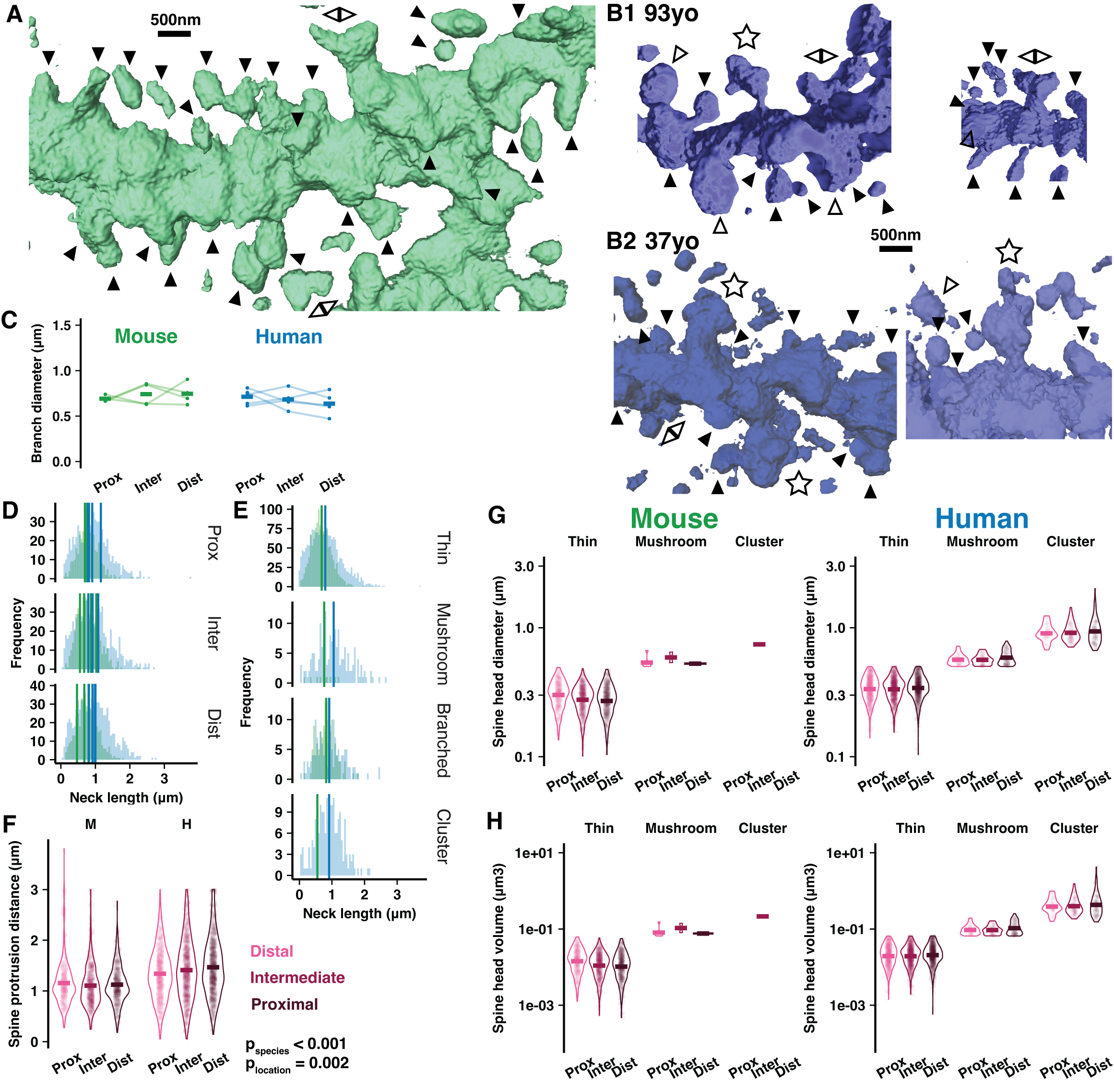

### Supplemental Figure 4

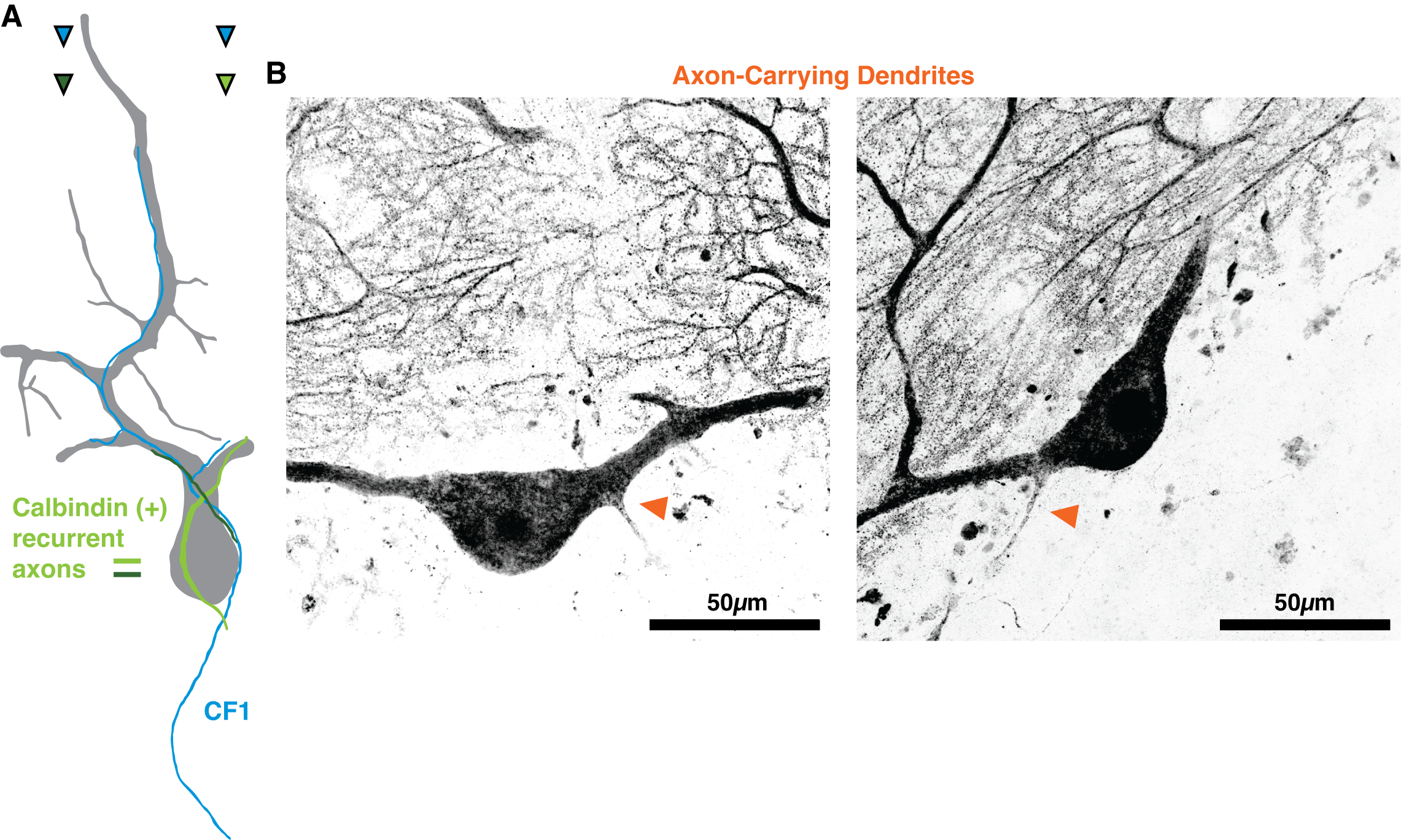

### Supplemental Figure 5

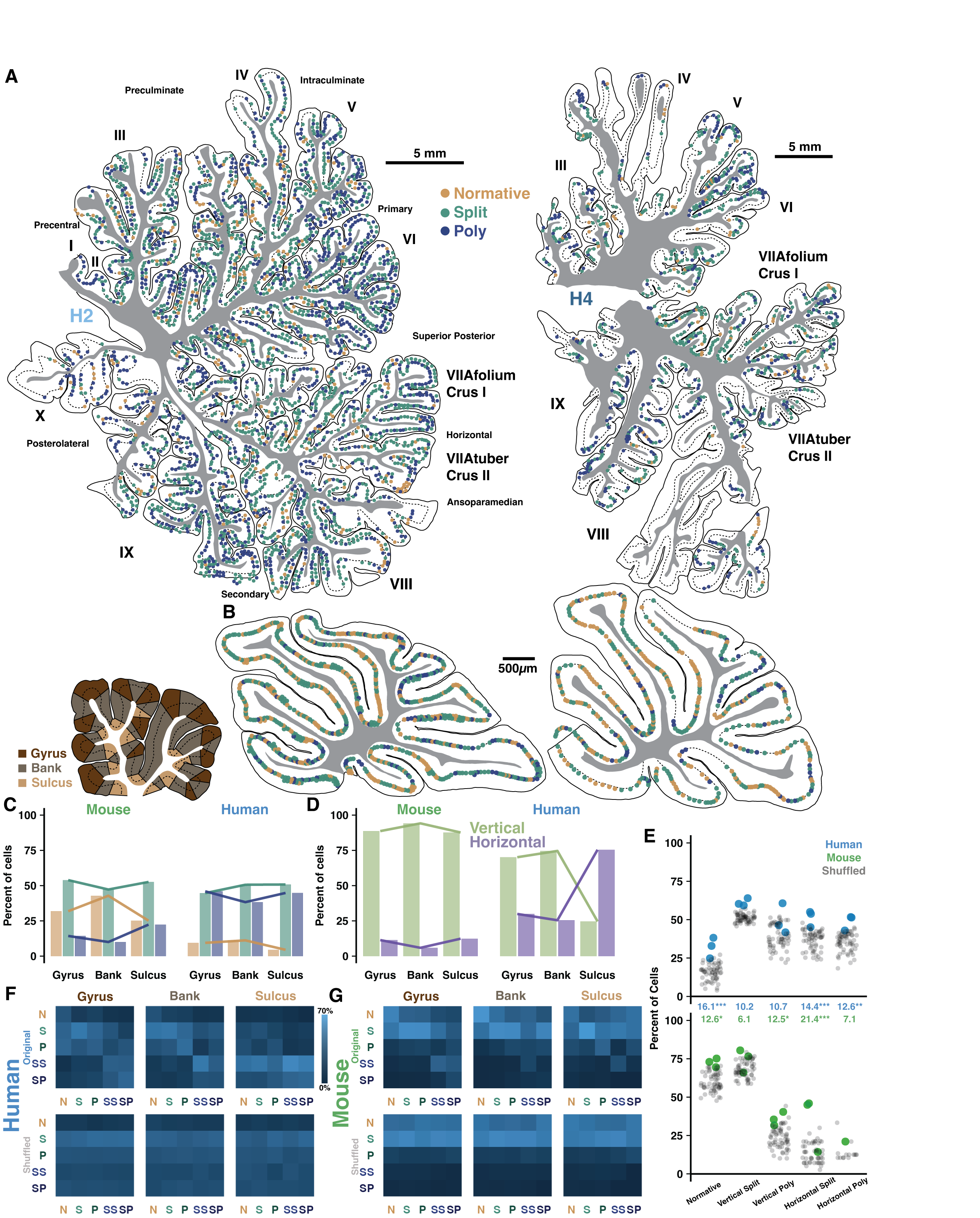

### Supplemental Figure 6

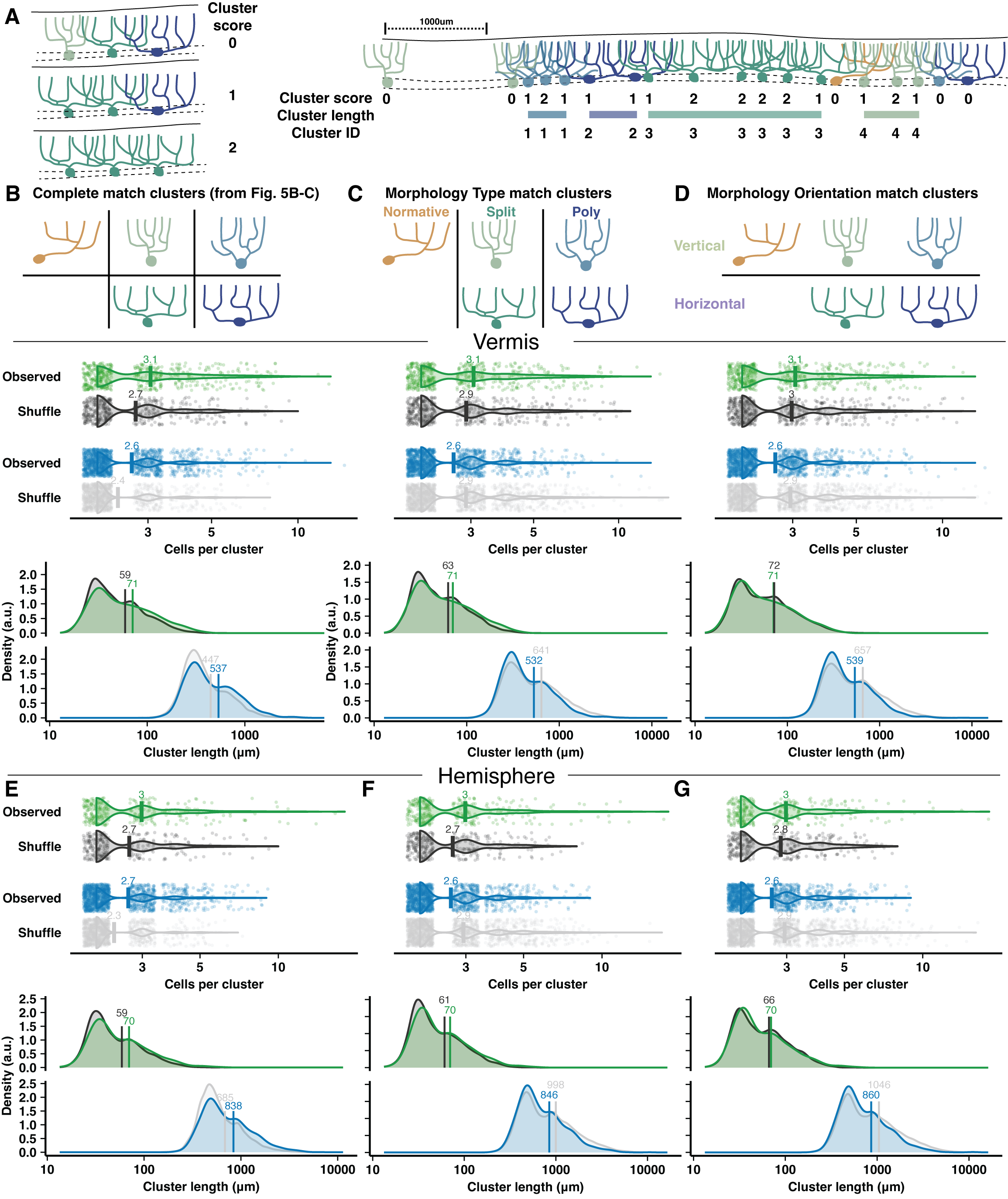

### Supplemental Figure 7

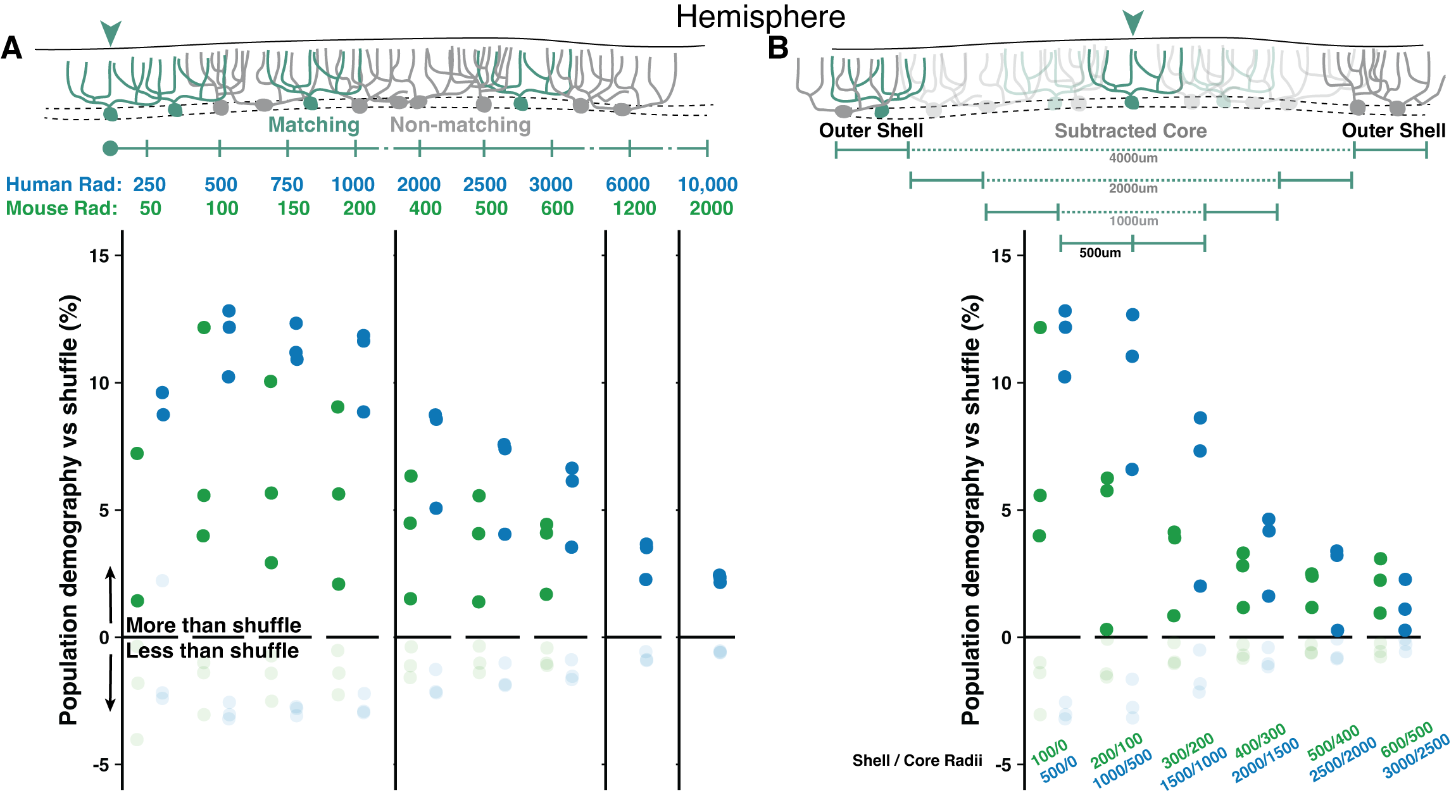

### Supplemental Figure 8

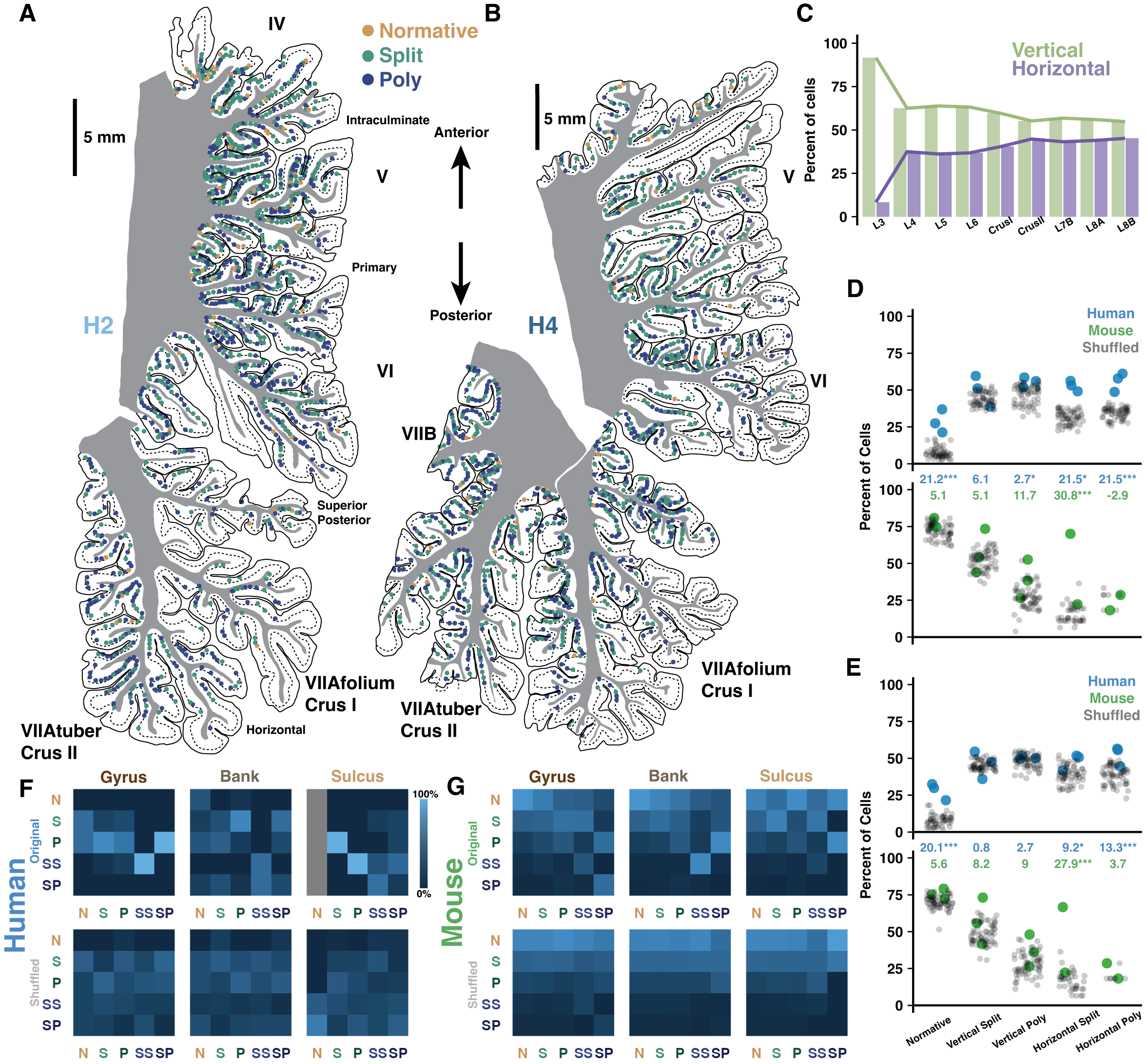

### Supplemental Figure 9

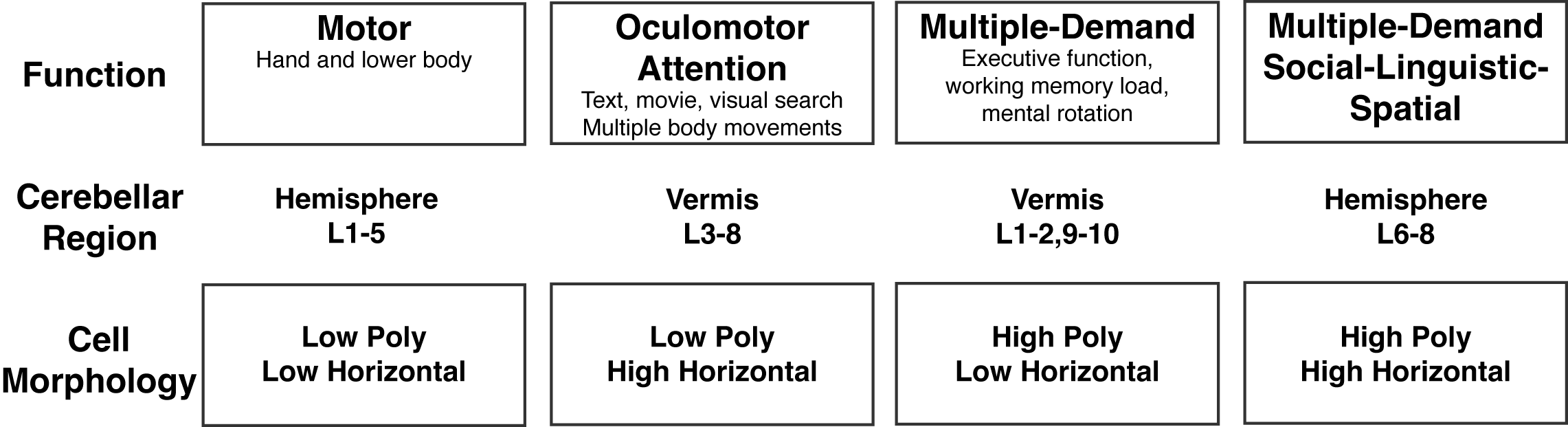
